# Radiation dermatitis in the hairless mouse model mimics human radiation dermatitis

**DOI:** 10.1101/2024.05.21.595074

**Authors:** Jessica Lawrence, Davis Seelig, Kimberly Demos-Davies, Clara Ferreira, Yanan Ren, Li Wang, Sk. Kayum Alam, Rendong Yang, Alonso Guedes, Angela Craig, Luke H. Hoeppner

## Abstract

Over half of all people diagnosed with cancer receive radiation therapy. Moderate to severe radiation dermatitis occurs in most human radiation patients, causing pain, aesthetic distress, and a negative impact on tumor control. No effective prevention or treatment for radiation dermatitis exists. The lack of well-characterized, clinically relevant animal models of human radiation dermatitis contributes to the absence of strategies to mitigate radiation dermatitis. Here, we establish and characterize a hairless SKH-1 mouse model of human radiation dermatitis by correlating temporal stages of clinical and pathological skin injury. We demonstrate that a single ionizing radiation treatment of 30 Gy using 6 MeV electrons induces severe clinical grade 3 peak toxicity at 12 days, defined by marked erythema, desquamation and partial ulceration, with resolution occurring by 25 days. Histopathology reveals that radiation-induced skin injury features temporally unique inflammatory changes. Upregulation of epidermal and dermal TGF-ß1 and COX-2 protein expression occurs at peak dermatitis, with sustained epidermal TGF-ß1 expression beyond resolution. Specific histopathological variables that remain substantially high at peak toxicity and early clinical resolution, including epidermal thickening, hyperkeratosis and dermal fibroplasia/fibrosis, serve as specific measurable parameters for in vivo interventional preclinical studies that seek to mitigate radiation-induced skin injury.

## Introduction

Radiation therapy is integral for control of many tumors and is prescribed for more than 50% of cancer patients in the United States ^[1–3]^. Approximately 95% of cancer patients receiving radiation therapy will develop moderate to severe radiation-induced dermatitis or “radiation burn”, with effects ranging from dry desquamation and erythema to moist desquamation and full thickness ulceration ^[4–9]^. Not only does it cause pain, anxiety, and disruption of quality of life during and following treatment, its severity correlates to the likelihood of the development of chronic effects like fibrosis, telangiectasia, ulceration, and necrosis ^[6,9]^. In some patients, radiation dermatitis is sufficiently severe to limit the therapeutic dose administered for tumor control or will lead to a break in treatment, which compromises local control and survival ^[8–12]^. Each day that radiation is delayed decreases tumor control and increases mortality, thus it is prudent for patients to remain on schedule without interruptions ^[13–15]^. With the combination of radiation therapy and targeted drugs like cetuximab that cause a skin rash, moderate to severe radiation dermatitis occurs at a higher incidence and for a longer duration compared to radiation therapy alone ^[16–19]^. Even with routine adoption of advanced radiotherapy techniques like intensity modulated radiation therapy (IMRT), radiation dermatitis remains common and causes treatment delays in up to 50% of patients ^[20,21]^.

There is no effective prevention or treatment for radiation dermatitis ^[7,11,22–31]^. The most widely adopted recommendation is for patients with dermatitis to keep the site clean using dilute soap and water and allow wound healing to occur ^[9,16,32]^. The molecular pathogenesis of radiation dermatitis is incompletely understood, and irradiated skin is rarely sampled repeatedly to evaluate signaling pathways. A major factor contributing to our limited molecular understanding is the lack of an optimized animal model of radiation dermatitis that provides comprehensive information regarding the spectrum of inflammation and pain that occurs. Animal models are essential for the advancement of novel agents targeted for use in the human cancer patient, thus it is prudent to have a well-defined model that translates favorably to the clinic. Previous rodent radiation dermatitis studies have suffered significant shortcomings, including variations in mouse strain, anatomical site and field size irradiated, radiation dosing details, radiation equipment, monitoring, and output measures ^[33–37]^. These factors contribute to an inability to effectively repeat, improve upon, or compare experimental approaches. Outbred SKH-1 mice are the most commonly used mouse strain for dermatologic studies ^[38]^ and have been widely used to study cutaneous effects of UVB irradiation ^[39–45]^. This strain represents an emerging model to evaluate dermatitis following ionizing radiation used for therapeutic purposes in oncology ^[33,46,47]^. Hairless mice are ideal because their wound healing has been well characterized, and their skin mimics human sebaceous skin most affected by radiation dermatitis ^[38,48]^. The objective of this study was therefore to characterize development of radiation-induced skin injury in an SKH-1 mouse model, highlighting distinct temporal stages of clinical and/or pathologic injury. The findings reported here provide a platform on which to objectively evaluate cellular signals contributing to these distinct stages, such that effective mitigators can be developed to reduce the severity, duration, and/or discomfort associated with radiation dermatitis.

## Results

### Radiation-induced clinical dermatitis occurs following 30 Gy irradiation

An initial pilot dose escalation study was performed to optimize the 30 Gy radiation dose used to induce dermatitis. SKH-1 mice were initially treated and evaluated for the development of acute dermatitis following a single dose of 15 Gy, 20 Gy, 25 Gy and 30 Gy (Supplementary Fig 1). A single treatment of 30 Gy using 6 MeV electrons was sufficient to induce severe (grade 3) toxicity 12 days (d) with resolution by 28d. A single fraction of 30 Gy radiation delivered with 6 MeV electrons induced severe clinical radiation skin injury (Fig. 1), defined by marked erythema, desquamation and partial ulceration. Radiation dermatitis grade significantly (p<0.0001) increased until peak toxicity with partial resolution by the end of the study period. Mean grade at peak toxicity on day 12 (2.56 ± 0.18) was significantly higher (p<0.0001) than all other timepoints evaluated (Table 1). Mean grade at resolution at day 22 (0.56 ± 0.24) was not significantly different than mean grade at baseline, 2 hours, and 5 days.

**Figure 1.**
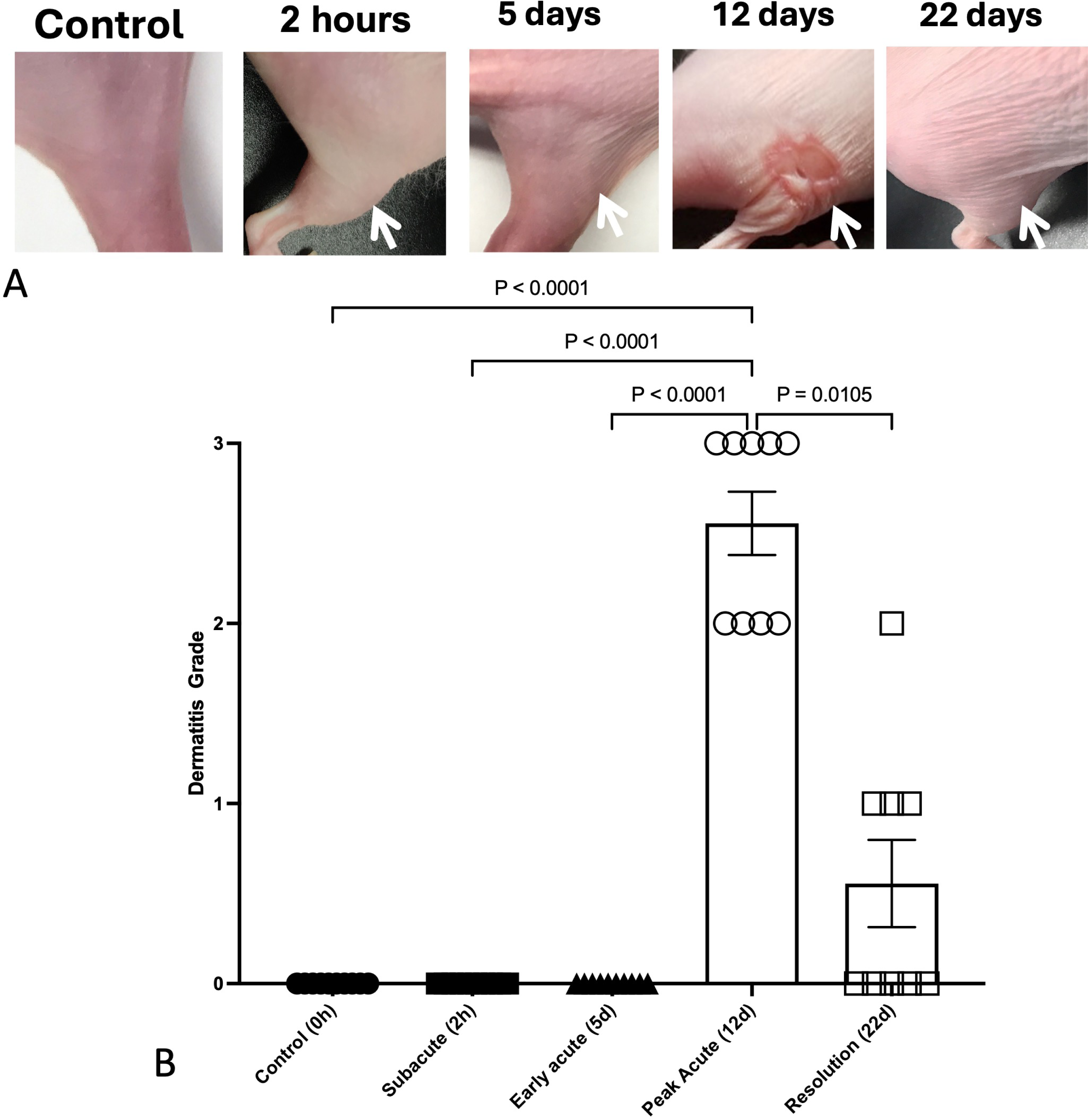
Temporal development of radiation-induced dermatitis in SKH-1 mice. Radiation dermatitis significantly (p < 0.0001) increases in severity following 30 Gy single fraction irradiation to the right hindlimb skin in 11-12 week old SKH-1 mice (N = 9-10/group). Representative photos of the right hindlimb (A) are shown following 30 Gy radiation delivered with 6 MeV electrons in comparison to unirradiated skin. Mean grade significantly increased at peak toxicity on day 12 (B) with partial resolution by day 22. Data are presented as the mean ± SEM at each defined timepoint following irradiation. Significant (p < 0.05) differences are shown following Kruskal-Wallis with Dunn’s multiple comparison analysis.

**Table 1.** Radiation dermatitis grade over time in 11-12 week old SKH-1 mice (N=9-10 per time point) following 30 Gy radiation to the skin of the right proximal hindlimb.

| Time | Mean grade ± SEM |
| --- | --- |
| 0h | 0.00 ± 0 |
| 2h | 0.00 ± 0 |
| 5d | 0.00 ± 0 |
| 12d | 2.56 ± 0.18 |
| 22d | 0.56 ± 0.24 |

### Temporally unique histopathologic skin inflammatory changes, including epidermal thickening, hyperkeratosis, and dermal fibroplasia/fibrosis contribute to radiation-induced injury

There were distinct histopathological changes over time with increasing total inflammatory score by day 12 that partially resolved by day 22 (Fig. 2). Histopathologic scores at 12 days and 22 days after irradiation were significantly higher than scores within unirradiated skin and within irradiated skin at 2 hours and 5 days (Table 2).

**Figure 2.**
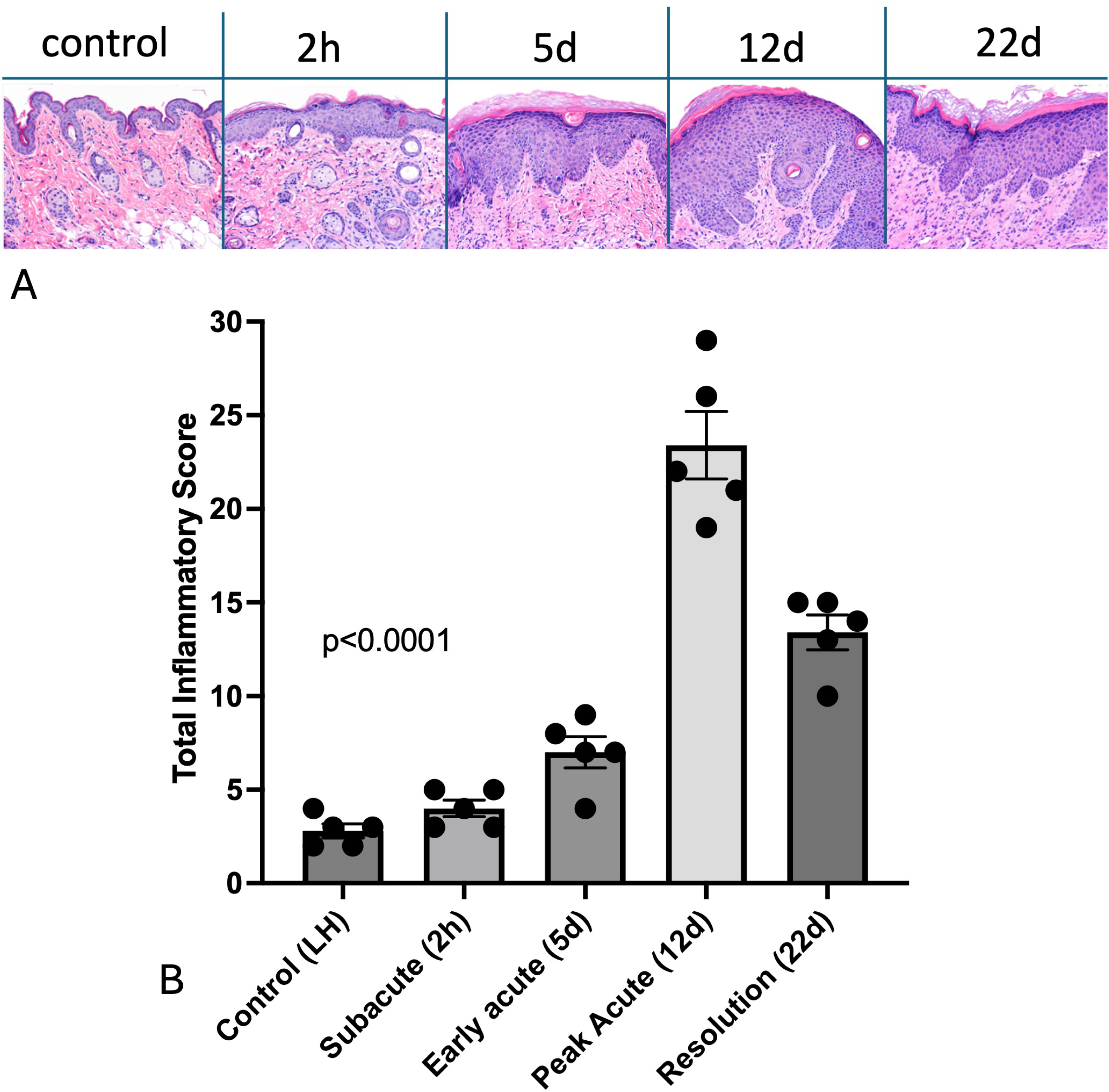
Radiation-induced dermatitis is characterized by measurable inflammatory changes over time. (A) Representative H&E images of radiation-induced skin pathology over time following irradiation 30 Gy single fraction irradiation prescribed to the skin of the right hindlimb/hip. (B) Total inflammatory score is represented as individual values and mean ± SEM for control skin from the left hindlimb (LH) and for irradiated skin from the right hindlimb (N=5 per time point) at designated time points following irradiation. The p value was calculated following one-way ANOVA.

**Table 2:** Significant differences in total inflammatory score in skin of SKH-1 mice at designated time points (N=5 per group) following 30 Gy radiation.

|  | 2h | 5d | 12d | 22d |
| --- | --- | --- | --- | --- |
| 0h | ns | ns | <0.0001 | <0.0001 |
| 2h |  | ns | <0.0001 | <0.0001 |
| 5d |  |  | <0.0001 | 0.0020 |
| 12d |  |  |  | <0.0001 |

*ns* represents “not significant” with *p* value > 0.05.

Significant changes in almost all histopathologic measures of inflammation were observed 12 days following irradiation, compared to control and subacute (2h) samples (Figure 3). Scores for epidermal ulceration, epidermal thickening, hyperkeratosis, glandular loss, dermal fibroplasia/fibrosis, dermal mononuclear, mastocytic, and neutrophilic inflammation, and hypodermal inflammation were significantly increased. Scores for hyperkeratosis (Fig. 3C), glandular loss (Fig. 3D), and dermal fibroplasia/fibrosis (Fig. 3E) remained significantly increased compared to unirradiated control 22d after irradiation. While specific inflammatory changes at 5d were not significantly different than control skin samples, increased glandular loss along with dermal mononuclear and neutrophilic inflammation were observed at 5d relative to unirradiated controls. Except for dermal pyogranulomatous inflammation, inflammatory scores significantly correlated to clinical grade (Table 3). Total inflammatory score, epidermal thickening, hyperkeratosis, and dermal fibroplasia/fibrosis strongly (r > 0.80) and positively correlated to clinical grade.

**Figure 3.**
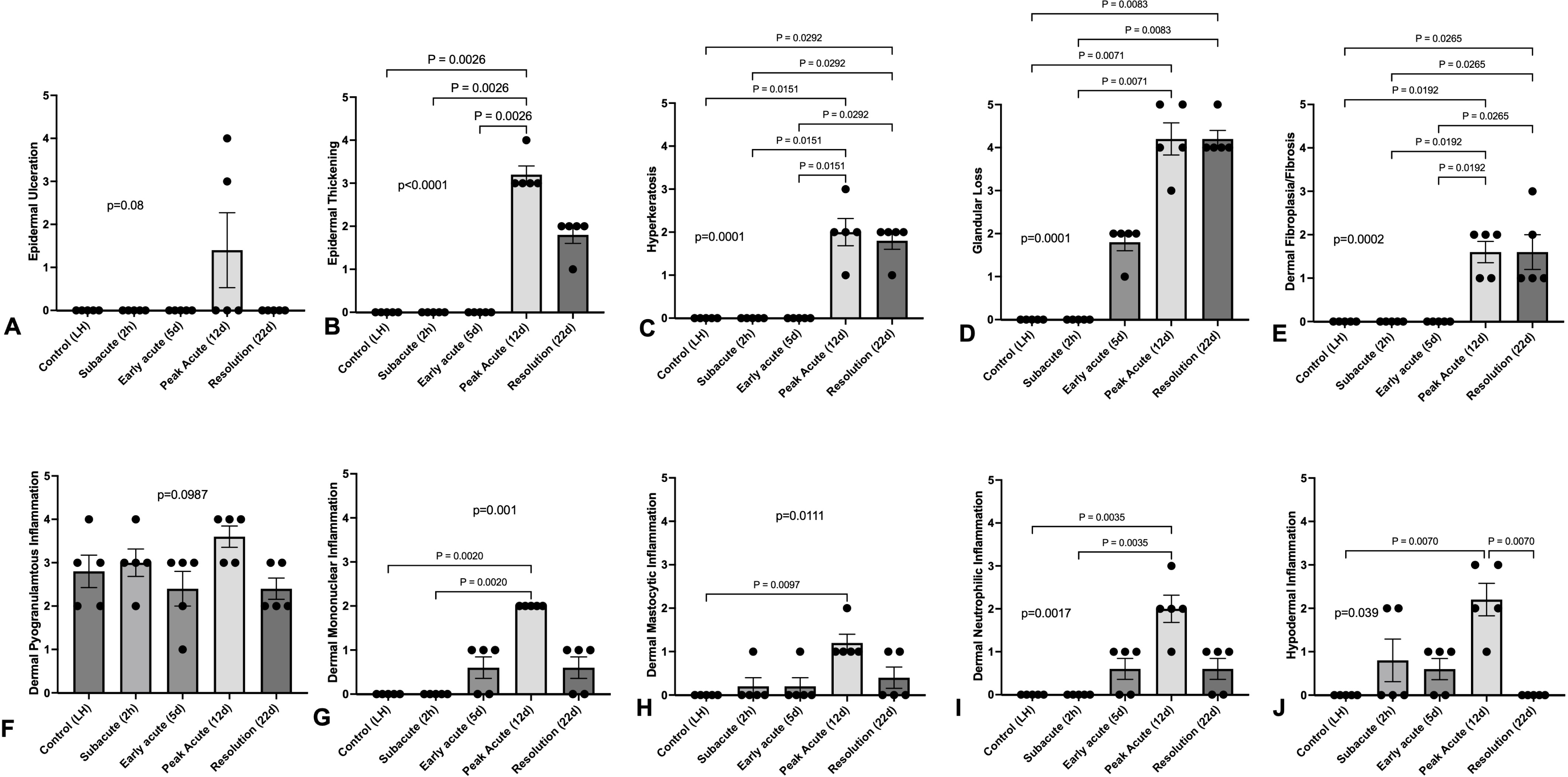
Radiation induced significant changes in most histopathologic measures of inflammation at the time of peak clinical toxicity on day 12. Mean scores for measured histopathologic variables, including epidermal ulceration (A), epidermal thickening (B), hyperkeratosis (C), glandular loss (D), dermal fibroplasia/fibrosis (E), dermal pyogranulomatous inflammation (F), dermal mononuclear inflammation (G), dermal mastocytic inflammation (H), dermal neutrophilic inflammation (I) and hypodermal inflammation (J), and shown from the unirradiated control left hindlimb (LH) skin and from irradiated skin over time. Data are presented as individual values and mean ± SEM, with p values in lowercase on the graph representing Kruskal-Wallis analysis. Comparative p values in uppercase between bars were calculated by performing Dunn’s multiple variable post-test analysis (N=5 per timepoint). Significant values between timepoints are highlighted with the bar; p values represent Kruskal-Wallis results while P values represent Dunn’s post-hoc results.

**Table 3:** Clinical dermatitis grade significantly (p<0.05) and positively correlated to the total inflammatory score as well as to most individual histopathological features assessed.

| Variable | *r<sub>s</sub>* | *p* value |
| --- | --- | --- |
| Total inflammatory score | 0.7988 | <0.0001 |
| Epidermal ulceration | 0.4605 | 0.0205 |
| Epidermal thickening | 0.9612 | <0.0001 |
| Hyperkeratosis | 0.8356 | <0.0001 |
| Glandular loss | 0.7636 | <0.0001 |
| Dermal fibroplasia/fibrosis | 0.8477 | <0.0001 |
| Dermal mononuclear inflammation | 0.7643 | <0.0001 |
| Dermal mastocytic inflammation | 0.6382 | 0.0006 |
| Dermal neutrophilic inflammation | 0.7303 | <0.0001 |
| Hypodermal inflammation | 0.5015 | 0.0107 |

### Increased epidermal and dermal TGF-ß1 and COX-2 protein expression occur at peak dermatitis, with sustained epidermal TGF-ß1 expression beyond clinical resolution of peak toxicity

TGF-ß1 plays roles in mediation and regulation of acute skin injury, cutaneous wound healing, and chronic fibrosis ^[49,50]^. Therefore, we evaluated epidermal and dermal TGF-ß1 expression in unirradiated (N=8) and irradiated (N=4-6 per time point) skin samples. Mean epidermal and dermal TGF-ß1 immunoreactivity scores significantly increased at day 12 compared to unirradiated samples (Fig. 4A-C). Increased epidermal TGF-ß1 protein expression was sustained until at least 22 days following irradiation (Fig. 4A). Mean dermal TGF-ß1 protein expression (3.75) at day 12 was similar to mean expression at day 22 (3.50). However, day 22 was not significantly different than mean control TGF-ß1 protein expression (p = 0.0506).

**Figure 4.**
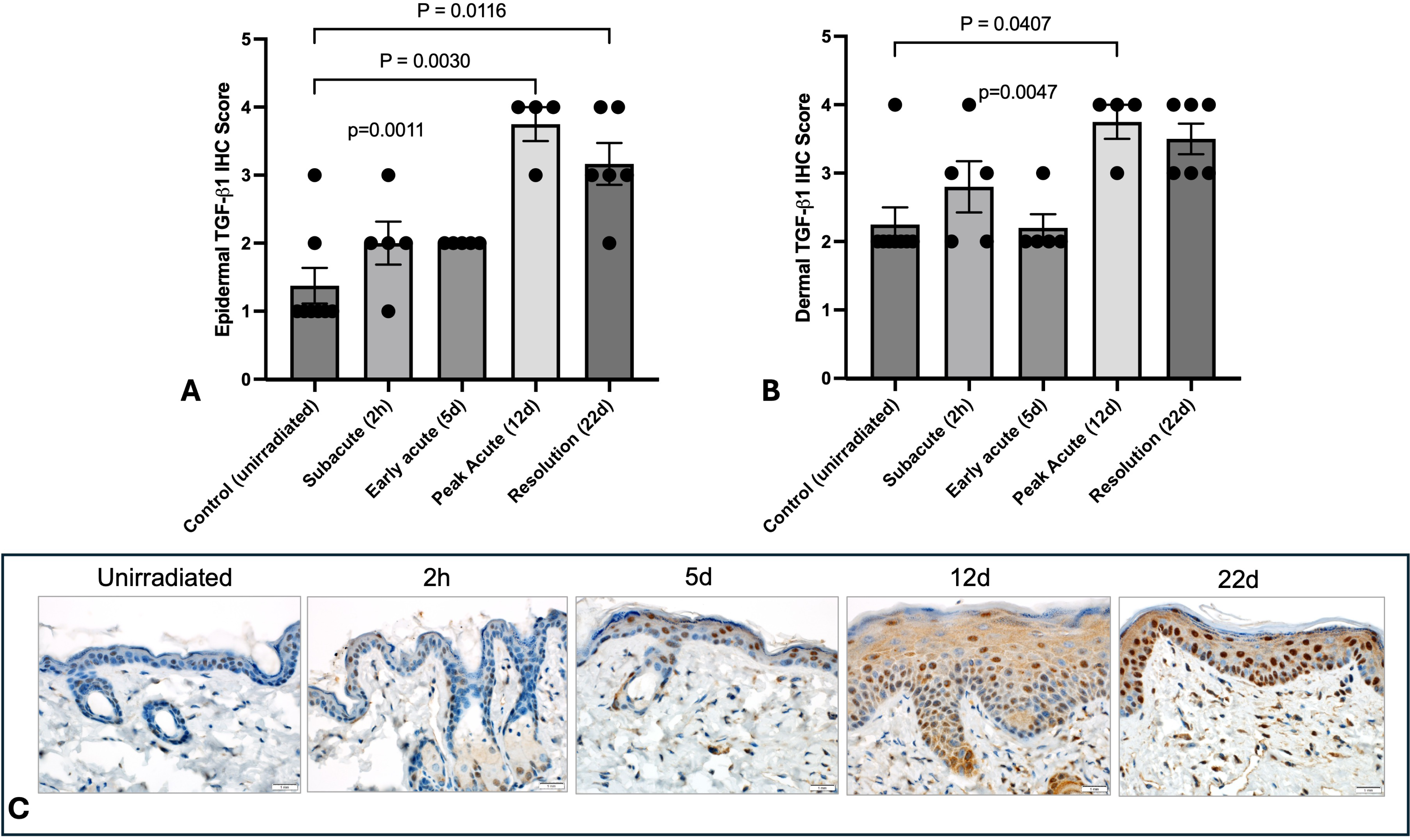
TGF-ß1immunoreactivity in irradiated skin from SKH-1 mice. Mean epidermal (A) and dermal (B) TGF-ß1immunoreactivity scores in unirradiated control skin (N=8) and irradiated skin (N=4-6) at designated time points after treatment. Data are presented as individual values and mean ± SEM, with p values in lowercase on the graph representing Kruskal-Wallis analysis. Comparative p values in uppercase between bars were calculated by performing Dunn’s multiple variable post-test analysis (N=5 per timepoint). Significant values between timepoints are highlighted with the bar; p values represent Kruskal-Wallis results while P values represent Dunn’s post-hoc results. (C) Representative tissue samples show normal positive TGF-ß1 immunohistochemical staining, represented as brown staining within the cellular cytoplasm, within the unirradiated dermis and epidermis. Progressively increased TGF-ß1 expression is demonstrated over time, with peak staining at day 12 and 22.

Because COX-2 plays a central role in a broad range of inflammatory processes in the skin, including hyperalgesia and edema ^[51]^, we evaluated COX-2 expression with the dermis and epidermis in irradiated (N=4-6 per time point) and control samples (N=6) (Fig. 5A-C). Epidermal and dermal COX-2 immunoreactivity scores were significantly higher at day 12 compared to unirradiated samples (Fig. 5A-B).

**Figure 5.**
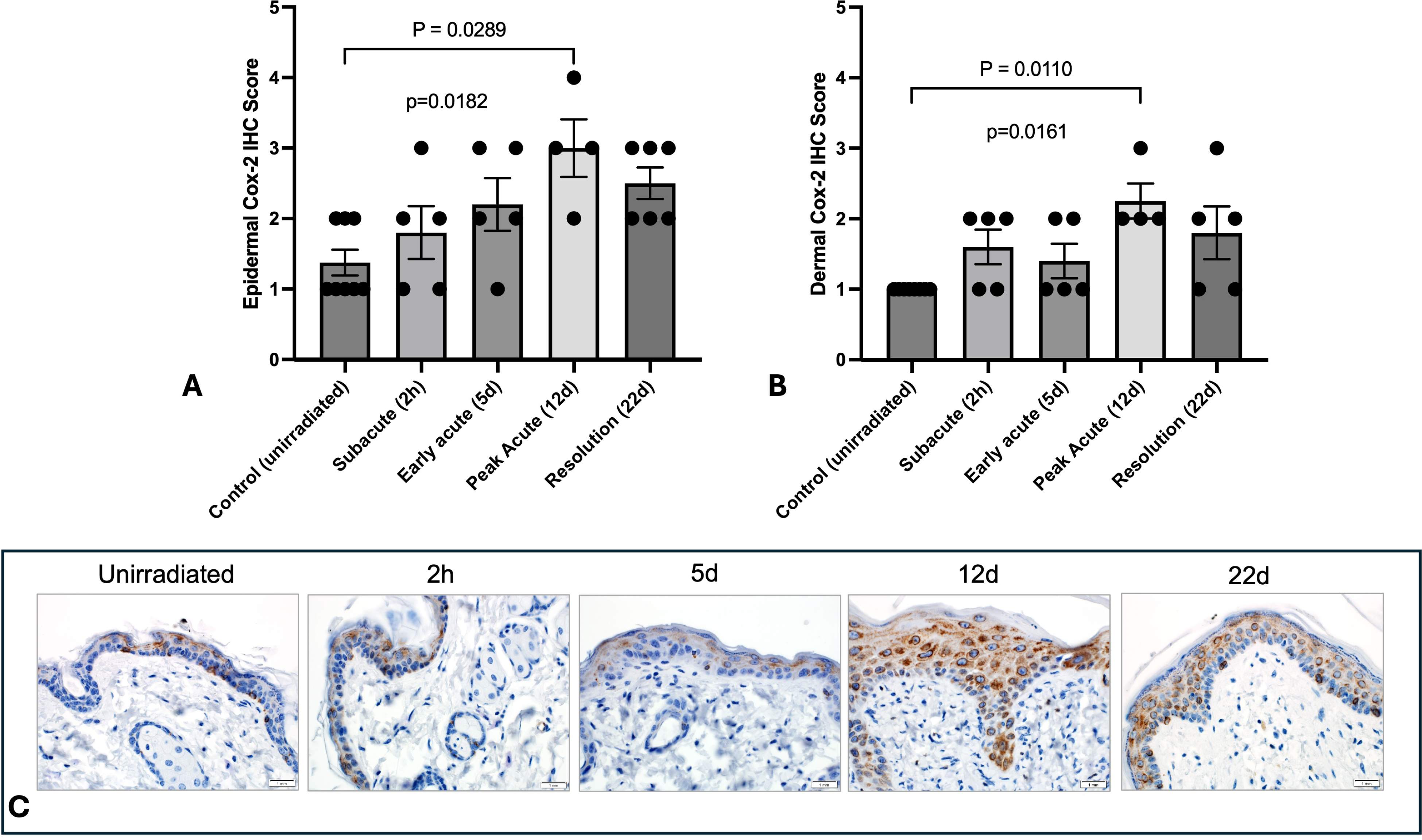
COX-2 immunoreactivity in irradiated skin from SKH-1 mice. Mean epidermal (A) and dermal (B) COX-2 immunoreactivity scores in unirradiated control skin (N=8) and irradiated skin (N=4-6) at defined time points. Data are presented as individual values and mean ± SEM, with p values in lowercase on the graph representing Kruskal-Wallis analysis. Comparative p values in uppercase between bars were calculated by performing Dunn’s multiple variable post-test analysis (N=5 per timepoint). Significant values between timepoints are highlighted with the bar; p values represent Kruskal-Wallis results while P values represent Dunn’s post-hoc results. (C) Representative tissue samples show normal positive COX-2 immunohistochemical staining, represented as brown staining within the cellular cytoplasm, within the unirradiated dermis and epidermis. Progressively increased COX-2 expression is demonstrated over time, with peak staining at day 12 and 22.

## Discussion

The hairless SKH-1 mouse strain is commonly used in translational dermatologic studies, and it is an ideal model for use in interventional studies where observations of inflammatory skin changes may be obscured by hair and pigment ^[33,38,46,48]^. The findings presented here describe a method of inducing robust radiation-induced dermatitis in the SKH-1 mouse, and the results highlight distinct temporal epidermal and dermal histopathologic changes that correlate to clinical radiation dermatitis grade. Clinical grading schemes, such as our modified CTCAE v5.0, provide a standardized approach to treatment-related adverse events and are important endpoint measures for studies that impact radiation-induced toxicity ^[52]^. Clinical grading is derived through standard, manual observations over time. The histopathologic changes that occur after irradiated skin are variable, may occur prior to clinical changes, and may be present despite apparent resolution. Understanding the specific inflammatory tissue changes underlying increasing clinical grade may improve the use and importance placed on clinical grade. In this study, nearly all histopathologic variables assessed significantly correlated to clinical dermatitis grade, which was highest at day 12 and 22. We examined a total inflammatory score that was comprised of discrete histopathologic insults to skin, as well as the individual histopathologic changes within the total score. Increased epidermal thickening, hyperkeratosis, and dermal fibroplasia/fibrosis most strongly correlated to increased clinical grade. Of these, hyperkeratosis and dermal fibroplasia/fibrosis were also significantly higher at day 12 and day 22 compared to unirradiated skin. Glandular deficiency was an additional histopathologic measure of radiation-induced injury that remained significantly high at day 22. Although not significantly different than baseline, early histopathologic changes were observed 5 days after irradiation, including glandular loss, monocytic and neutrophilic inflammation.

The skin barrier acts as a critical protector for the body against external environmental hazards. Recent data has shown that up to 66% of cancer patients have at least one non-cancer related co-morbidity, while 50% have multiple co-morbidities ^[53]^, with the highest prevalence in lower socio-economic groups ^[53–55]^. Common co-morbidities like hypertension, diabetes and heart disease are associated with unique skin conditions that affect skin integrity and healing ^[56,57]^. Maintenance of skin integrity and barrier function is therefore of incredible importance, because the skin is the first line of protection against microbes, toxins, sunlight and other external exposures ^[58]^. Glandular loss has been recently shown to impair the skin barrier in SKH-1 mice treated with 20-40 Gy to the hindlimb ^[46,47]^. Glandular loss was seen as early as 4 days following 40 Gy, and within 6 days following 20-30 Gy ^[46]^. Our findings of glandular loss prior to clinical dermatitis, although not significantly different than unirradiated skin when assessed by our 5-point histopathologic scale, align with these prior reports. Importantly, our data show that glandular loss develops early and remains significantly impacted beyond initial clinical recovery of dermatitis. Potential therapeutic interventions that preserve dermal sebaceous glands during and after irradiation may improve skin integrity and promote healing. Clinical grade moderately correlated with glandular loss, and additional measures may be beneficial for amelioration of pre-clinical changes.

Increasing epidermal thickening and hyperkeratosis were strongly correlated to increasing clinical grade in our SKH-1 model. These epidermal changes occur secondary to the inflammatory cascade that occurs following irradiation, and they are well recognized in the irradiated skin of cancer patients ^[6,8]^. Epidermal damage and structural keratin changes, together with the preceding inflammatory signals, disrupt the skin barrier and can foster dysbiosis and chronic inflammation that further perpetuates skin injury ^[59]^. Interestingly, a recent study identified early epidermal thickening and hyperkeratosis in human skin-equivalent tissue models following single fractions of either 2 Gy or 10 Gy delivered with 6 MeV electrons, similar to our radiation delivery method ^[60]^. This study utilized optical coherence tomography with subsequent histology to confirm visual findings and may provide a non-invasive means of measuring these two features in future radiation studies seeking to mitigate radiation dermatitis.

Because TGF-ß1 is a key mediator of tissue repair following injury and subsequent tissue fibrosis after irradiation, studies have suggested considering TGF-ß1 in the evaluation of early radiation dermatitis and its healing process ^[61–63]^. In our SKH-1 radiation dermatitis model, dermal fibroplasia and fibrosis strongly correlated to clinical grade. Our data also demonstrated an early increase in TGF-ß1 expression within the epidermis and dermis, with sustained high expression in the former at the end of the study period, when the mean clinical dermatitis grade was close to baseline (mean 0.56). This sustained TGF-ß1 expression in SKH-1 mice is similar to prior reports in other murine strains to describe early, robust and long-term TGF-ß1 mRNA expression following 50 Gy irradiation ^[64,65]^. This also mirrors data in humans demonstrating significantly upregulated TGF-ß1 mRNA expression following preoperative radiation treatment ^[66]^. TGF-ß1 signaling regulates wound repair in irradiated skin by stimulating fibroblast, neutrophil and macrophage infiltration, which our 5 day histopathologic data support. Of note, recent studies have shown that Smad3-null mice have faster healing, reduced inflammation, and reduced early scarring within irradiated skin compared to mice with intact Smad3 ^[62,67]^. Smad3 is a critical downstream mediator of TGF-ß1 signaling that mediates several repair processes in skin, including inflammation, induction of epithelial-to-mesenchymal transdifferentiation, keratinocyte migration, and granulation tissue formation ^[62,63,67]^. Data supports use of SKH-1 mice as an appealing model for preclinical investigation of inhibitors of TGF-ß1-Smad3 signaling to reduce dermatitis severity and duration. Inhibiting this pathway may have dual benefit, as TGF-ß1-Smad3 signaling is implicate in the induction and maintenance of therapeutic resistance for some breast cancers ^[68]^.

The role of COX-2 within the skin and its inflammatory responses is varied. COX-2 has received attention as a therapeutic target by which to mitigate dermatitis in the past because it is pro-inflammatory, pro-angiogenic and associated with pain ^[69]^. Several studies investigating SKH-1 mice have highlighted that COX-2 mediates UVB-irradiation induced inflammatory responses in the skin ^[39–42,70]^. One study reported reduced skin damage following irradiation in female C3H/He mice following treatment with a highly selective COX-2 inhibitor ^[71]^. A randomized controlled trial failed to demonstrate reduced radiation dermatitis with the use of highly COX-2 selective drugs ^[72]^. Increased epidermal and dermal COX-2 expression was noted at the time of severe dermatitis (day 12) compared to unirradiated skin samples. It is important to note that mice evaluated in our studies received 1-4 doses of carprofen, a COX-2 inhibitor, beginning on day 11 or 12 once daily to reduce lameness and pain associated with limb dermatitis. This was an ethical decision that aligned with our institutional policies to maintain animal welfare. It is possible that epidermal and dermal COX-2 protein expression from the day 12 and day 22 skin samples were dampened by systemic administration of carprofen, and expression may have been higher in its absence. Because COX-2 mediates a host of pro-inflammatory and pro-nociceptive signals, evaluation of COX-2 within SKH-1 skin in future studies may be beneficial.

Radiation therapy is prescribed for more than 50% of the 1.8 million cancer patients in the US ^[1,2,73,74]^. Clinical signs of radiation dermatitis range from dry desquamation and erythema to moist desquamation and full thickness ulceration ^[4–9]^. Its severity correlates with chronic effects like fibrosis, telangiectasia, hyperpigmentation, and necrosis ^[6,9]^. Acute dermatitis causes pain and anxiety, while disrupting quality of life ^[75]^. In people of color, the severity of acute dermatitis ^[76]^ and the impact of chronic skin changes like hyperpigmentation are particularly detrimental to quality of life ^[77]^. Severe radiation dermatitis leads to cancer treatment interruptions in some patients, which significantly reduces tumor control and survival ^[8–15]^. Despite technological advances, such as intensity modulated radiation therapy, dermatitis causes treatment delays in up to 50% of patients ^[20,21]^. Management of radiation dermatitis is costly and often requires specialty symptom management due to skin effects ^[78,79]^. Data suggests that nursing encounters, cost of wound care consumables, and direct nursing costs could all be significantly reduced with implementation of strategies to reduce acute radiation skin toxicities ^[80]^. There is no effective prevention or treatment for radiation dermatitis ^[7,22–31]^. Despite prior studies, the most widely adopted recommendation is to keep irradiated skin clean and allow wound healing to occur ^[9,16,32,81,82]^.

Our data describing radiation-induced skin injury in the SKH-1 model is also useful beyond the context of therapeutic exposures. Cutaneous injuries can develop in normal human skin following a wide variety of radiation exposures, including nuclear device fallout, nuclear energy accidents, nuclear testing, medical exposures, and industrial overexposures ^[83–86]^. Indeed, skin damage is the most common radiation injury in humans ^[87]^. Important lessons from victims of nuclear disasters (i.e., atomic bombings of Hiroshima and Nagasaki, the Chernobyl nuclear accident) highlight the array of clinical manifestations of skin injury, the lack of effective treatment or pain management for resulting dermatitis, and the negative impact of injured skin on the likelihood of fatal systemic complications ^[88,89]^. Additionally, medical exposures can result from diagnostic procedures or therapeutic exposures for treatment ^[79,85,90,91]^. Over one million cases of diagnostic fluoroscopy-guided interventions occur annually in the US, and the frequency of complex interventional procedures that require longer radiation exposures have increased ^[92]^.

There is a clear clinical need to effectively mitigate radiation dermatitis, with implications for human health beyond medical and therapeutic radiation exposures. Our studies support the inclusion of the SKH-1 mouse as a preclinical model for radiation-induced dermatitis, as histopathologic features like glandular loss and TGF-ß1 protein expression may serve as endpoint measures following intervention. Clinical dermatitis grading in SKH-1 mice correlates well to histopathologic variables associated with epidermal and dermal injury. Specific histopathologic measures that remained significantly high at peak toxicity and at early resolution, namely epidermal thickening, hyperkeratosis and dermal fibroplasia/fibrosis, may be used to as distinct target variables to evaluate in future studies using SKH-1 mice to mitigate radiation-induced skin injury.

## Methods

### Mice

11-12 week old female SKH-1 mice were purchased (Charles River Laboratories) and used for all experiments. All experiments were approved by and performed in accordance with the Institutional Animal Care and Use Committee (IACUC Protocol #1808-36331A). Mice were housed in a group of 4 or 5 animals and were randomly assigned to housing upon arrival at the institution by Research Animal Resources staff. Mice were euthanized by carbon dioxide proceeded by exsanguination following the Institution’s IACUC Criteria for Carbon Dioxide Euthanasia Guidelines.

### Radiation

Mice were immobilized with ketamine (100 mg/kg) and xylazine (2 mg/kg) administered intraperitoneally 2-5 minutes prior to irradiation. All anesthetic events were overseen or carried out by a veterinarian with laboratory animal expertise. An initial pilot dose escalation study was performed to determine the target radiation dose to induce significant grade 3 dermatitis. Upon heavy sedation, mice were treated with a single dose of 15 Gy, 20 Gy, 25 Gy or 30 Gy. Following determination of 30 Gy as the target dose for all experiments, mice were treated with 30 Gy radiation to the skin surface using 6 MeV electrons with a custom 2 x 2 cm cutout (Varian iX, Varian Medical Systems, Inc, Palo Alto CA). Skin over the right proximal hindlimb was targeted in all mice. Skin over the left hindlimb served as a control. Tissue equivalent bolus (1 cm) was placed on the surface of the skin to provide sufficient dose build-up to the level of the skin with source-to-surface distance of 100 cm. Dose delivered to irradiated (right hindlimb) and unirradiated skin (left hindlimb) was verified via radiochromic film dosimetry (GAFchromic™) to ensure the dose was delivered as prescribed. ^[93]^ Following irradiation, dermatitis was graded daily using a modified Common Toxicity Criteria for Adverse Events (CTCAE v5.0) (Supplementary Table 1) ^[52]^. Because dermatitis was associated with pain and lameness, and pain was not an endpoint tested, we adhered to our institutional policy to maintain animal welfare and mice were treated with subcutaneous (Zoetis, Kalamazoo, MI) at a dosage of 5 mg/kg every 24 hours for 1-4 days beginning on day 11-12.

### Histopathology

To characterize pathologic changes over time, skin and subcutaneous histopathology were evaluated in unirradiated (control) skin from the left hindlimb and at 2 hours (h), 5 days (d), 12d, and 22d post irradiation. These time points were considered representative of acute injury (2h) early induction (5d), peak toxicity (12d), and initial resolution (22d) of radiation dermatitis. Skin from unirradiated and irradiated sites from each mouse was collected immediately following euthanasia. Skin was fixed in 10% neutral buffered formalin for 24h and subsequently embedded in paraffin wax. Four-micron tissue sections were deparaffinized in xylene and subsequently rehydrated in graded alcohol. Slides were stained with Harris Modified Hematoxylin with acetic acid (EXPREDIA, Kalamazoo, MI, Cat# 7221). The slides were dipped first into acid water (0.15% HCL, Acros Organics, Fair Lawn, Cat# NJAC124210010), followed by running tap water, and finally in ammonium water (2.8% of ammonium hydroxide 28-30%, Newcomer Supply, Middleton, WI, Cat# 1006A). The slides were counterstained with Eosin (Leica Biosystems, Deer Park, IL, Cat# 3801600). The slides were dehydrated in graded alcohol and xylene before coverslip-mounted using permount mounting media (Leica Biosystems, Deer Park, IL, Cat# 3801731). Hematoxylin and eosin (H&E)-stained sections were evaluated by a board-certified veterinary pathologist [American College of Veterinary Pathologists (ACVP)] on a 5-point scale for epidermal ulceration, epidermal thickening, hyperkeratosis, glandular loss, dermal fibrosis / fibroplasia, dermal inflammation (including pyogranulomatous inflammation, monocytic inflammation, mastocytic inflammation, and neutrophilic inflammation) and hypodermal inflammation according to a modified version of a previous publication ^[94]^. For each parameter, severity was defined as: 1 = minimally detectible, 2 = mild, 3 = moderate, 4 = marked and 5 = severe. A total inflammatory score comprised the sum of each histopathologic parameter score, with a maximum score of 50. Treatment-associated dermal inflammation was considered against the background of strain-associated follicle-centric inflammation in the control, unirradiated skin samples.

### Immunohistochemistry

To further characterize inflammatory pathways activated after radiation, sections of irradiated and unirradiated skin were immunostained for cyclooxygenase-2 (COX-2) and transforming growth factor-ß1 (TGF-ß1). For both COX-2 and TGF-B1 IHC, 4 μm formalin-fixed, paraffin-embedded tissue sections were deparaffinized and rehydrated, followed by antigen retrieval using either a high pH EDTA solution (COX-2) or a low pH citrate buffer (TGF-B1). After quenching endogenous peroxidase, immunohistochemistry was performed using one of two rabbit polyclonal primary antibodies (COX-2, Biocare, CRM-306 and TGF-B1, Invitrogen, PA1-29032) that were incubated for 30 minutes at room temperature. The antibodies were diluted at 1:200 and 1:100, respectively. Antibody binding was detected using the Rabbit Envision (Dako) secondary antibody kit. Diaminobenzidine was used as the chromogen and Mayer’s Hematoxylin (Dako) was used as the counterstain. Primary antibodies were substituted with appropriate negative control IgG for negative control slides. Samples were evaluated by a single pathologist and given a quantitative immunoreactivity score based on percentage of keratinocytes (epidermal samples) or nucleated cells (dermal samples) staining positive. Immunoreactivity scores were defined as: 0 = no staining detected, 1 = 0-25% cells (keratinocytes or nucleated cells) with positive immunostaining, 2 = 26-50% of cells with positive immunostaining, 3 = 51-75% of cells with positive immunostaining, and 4 = 76-100% of cells with positive immunostaining.

### Statistical Analysis

Commercially available software (Prism v10; GraphPad Software, Inc., San Diego CA) was used to evaluate data. Descriptive data was reported as mean ± standard error of the mean (SEM). Grade and histopathological variables were assessed for normality differences in grade and histopathological variables over time was determined using one-way analysis of variance (ANOVA) with Tukey’s multiple comparison test or Kruskal Wallis with Dunn’s multiple comparison test. Correlations between grade and histopathological variables were assessed using Spearman rank-order correlation coefficient (r_s_). Correlations were categorized as strong if r_s_=0.8-1.0, moderate if r_s_=0.4-0.8 and weak if r_s_=0.1-0.4. Statistical significance was set at p<0.05.

## Acknowledgements

This work was supported by funding from the Department of Veterinary Clinical Sciences and The Hormel Institute at the University of Minnesota. The authors would like to thank Jessica Coffey and Amy Morgan for their support for data acquisition.

## Author contributions

Conceptualization and design: JL, DS, LH

Development of methodology: JL, DS, AC, CF, AG, LH

Data acquisition: JL, DS, AC, CF, KDD

Data analysis: JL, DS, AC, CF, YR, LW, SKA, RY, LH

Writing and primary revision: JL, DS, AG, LH

Review & secondary revision: all authors (JL, DS, AC, CF, KDD, AG, YR, LW, SKA, RY, LH) read, provided secondary revision, and approved the submitted manuscript.

## Data availability statement

All data generated or analyzed during this study are included in this published article and its Supplementary Information files.

