## Supplementary Table 1 & Figure 1 for "Radiation dermatitis in the hairless mouse model mimics human radiation dermatitis"

**Supplementary Data**

Supplementary Table 1: Radiation-induced dermatitis grading scheme.

| Grade | Description |
|---|---|
| 0 | No change over baseline |
| 1 | Follicular, faint or dull erythema<br>Epilation<br>Patchy dry desquamation |
| 2 | Tender or bright erythema<br>Diffuse dry desquamation<br>Mild-moderate edema<br>Patchy non-dry desquamation |
| 3 | Diffuse non-dry glistening desquamation<br>Diffuse edema |
| 4 | Ulceration<br>Hemorrhage<br>Necrosis |
| 5 | Death due to dermatitis |

**Supplementary Figure 1.**

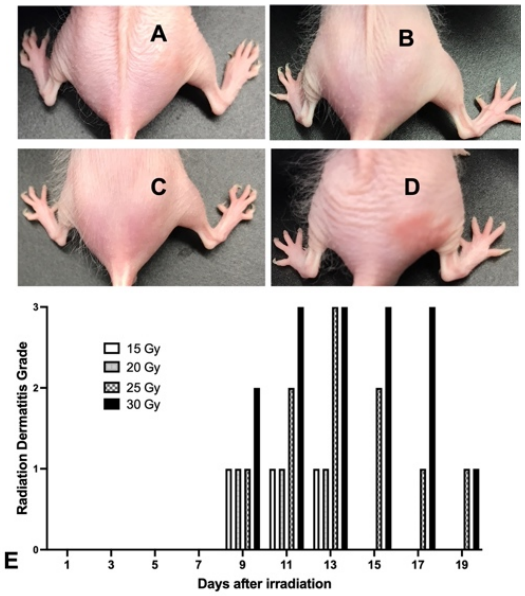


Radiation dermatitis in SKH-1 mice following pilot dose escalation study. SKH-1 mice (N=1 per dose) received one fraction of radiation at increasing doses directed to the skin over the right proximal hip. Photographs illustrating the radiation target site at peak toxicity following 15 Gy (A), 20 Gy (B), 25 Gy (C) and 30 Gy (D). Dermatitis grade was documented following treatment (E) to determine the radiation dose to be used in larger animal studies to produce persistent grade 3 dermatitis.
